# Non-monotonic fibril surface occlusion by GFP tags from coarse-grained molecular simulations

**DOI:** 10.1101/2021.10.21.465309

**Authors:** Julian C. Shillcock, Janna Hastings, Nathan Riguet, Hilal Lashuel

## Abstract

The pathological growth of amyloid fibrils in neurons underlies the progression of neurodegenerative diseases including Alzheimer’s and Parkinson’s disease. Fibrils form when soluble monomers oligomerise in the cytoplasm. Their subsequent growth occurs via nucleated polymerization mechanisms involving the free ends of the fibrils augmented by secondary nucleation of new oligomers at their surface. Amyloid fibrils possess a complex interactome with diffusing cytoplasmic proteins that regulates many aspects of their growth, seeding capacity, biochemical activity and transition to pathological inclusions in diseased brains. Changes to their surface are also expected to modify their interactome, pathogenicity and spreading in the brain. Many assays visualise fibril formation, growth and inclusion formation by decorating monomeric proteins with fluorescent tags such as GFP. Recent studies from our group suggest that tags with sizes comparable to the fibril radius may modify the fibril surface accessibility and thus their PTM pattern, interactome and ability to form inclusions. Using coarse-grained molecular simulations of a single alpha synuclein fibril tagged with GFP we find that thermal fluctuations of the tags create a non-monotonic, size-dependent sieve around the fibril that perturbs its interactome with diffusing species. Our results indicate that experiments using tagged and untagged monomers to study the growth and interactome of fibrils should be compared with caution, and the confounding effects of the tags are more complex than a reduction in surface accessibility. The prevalence of fluorescent tags in amyloid fibril growth experiments suggests this has implications beyond the specific alpha synuclein fibrils we model here.

## Introduction

Amyloid fibrils formed of misfolded proteins underlie many neurodegenerative diseases.[1] They are a component of Lewy bodies present in Parkinson’s disease,[2-4] amyloid plaques and neurofibrillary tangles in Alzheimer’s disease and cellular inclusions formed in Huntington’s disease among others.[5] The composition of these inclusions is complex and involves not only fibrils, but also lipids, membranous organelles, and other proteins.[4, 6] Fibrils may interfere with a cell’s homeostasis by their presence as rigid bodies in the cell and as the result of their complex interactome with proteins,[7] and cellular membranes and organelles[3] or their ability to activate cell death pathways and biochemical reactions occurring at specific surface domains or interaction hubs. Within pathological inclusions, the fibrils are typically heavily modified by enzymes that access their surface or the flexible termini of monomeric subunits that project into the surroundings (e.g., N- and C-terminal domains of aSyn). Post-translational modifications also influence fibril structure and surface properties[8-11] as well as their propensity to self-associate.[12]

Fibrils are nucleated from soluble monomeric proteins via two distinct mechanisms: 1) In the primary mechanism, short oligomers form in bulk solution and elongate by addition of monomers at their ends; 2) Secondary nucleation occurs when oligomers form at the surface of existing fibrils from monomers that have diffused onto them (Figure 1). These oligomers may then form branches from the existing fibril or detach and grow by the primary mechanism into new fibrils. Reaction rate models predict that secondary nucleation is a major contributor to fibril elongation.[13, 14] The fibril surface is also an important target for drugs to inhibit pathological amyloid fibrils as it contains multiple domains or binding sites for small molecules.[15] Access to the fibril surface is therefore a key determinant of pathological fibril growth, inclusion formation and potential therapeutic interventions. The ability of other proteins to modify or interact with the surface depends on the ease with which they can access it via diffusion.

**Figure 1.**
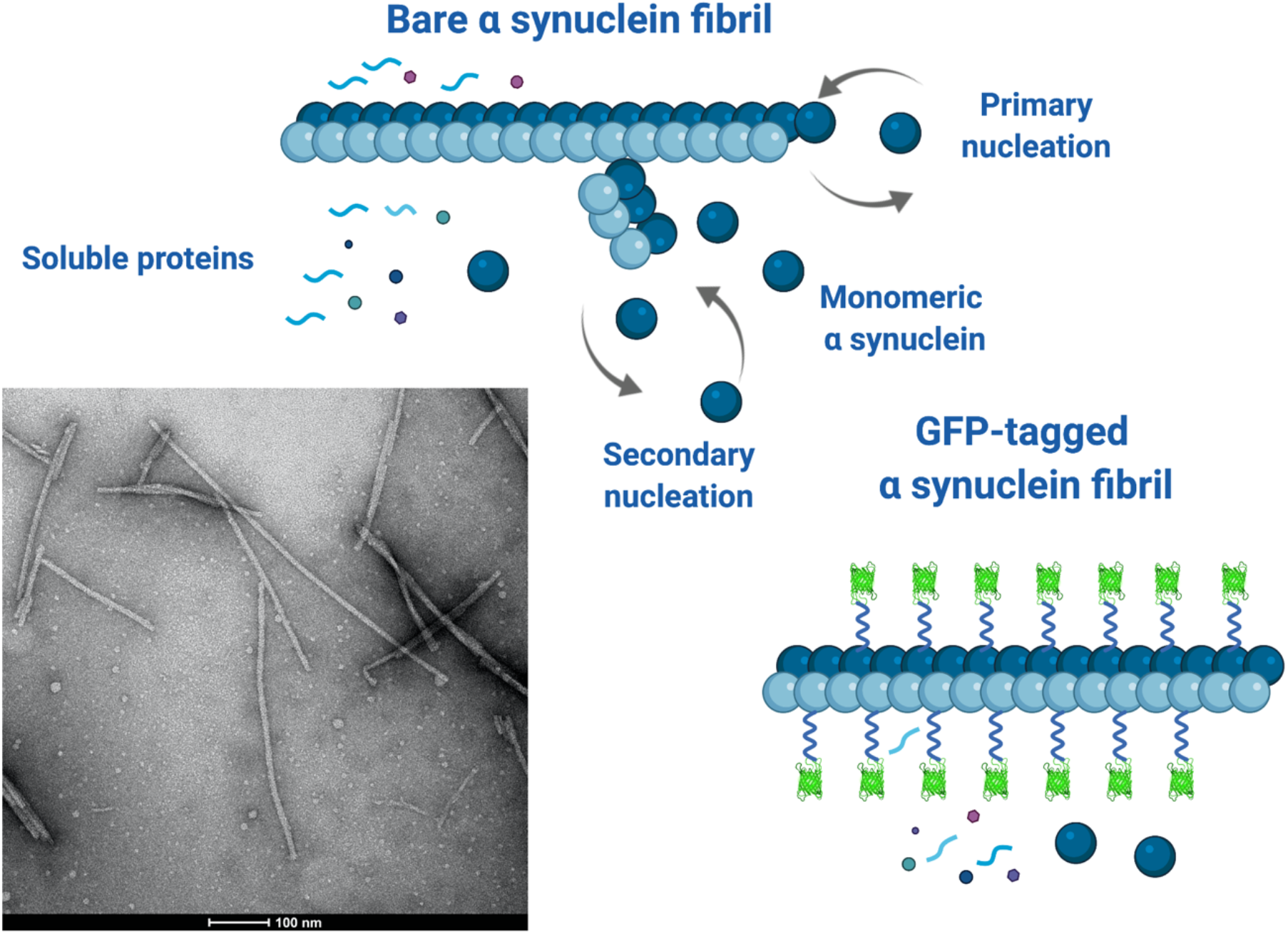
Cartoon of a bare aSyn fibril illustrating the primary and secondary nucleation mechanisms by which it elongates and forms new fibrils at its surface respectively. Soluble proteins and aSyn monomers/oligomers can diffuse to, and interact with, the surface of the fibrils giving rise to a dynamic interactome. The GFP-decorated fibril is shown undergoing less surface-mediated interactions due the occluding effects of the tags that can modify the fibril’s interactome. The EM image shows several bare aSyn fibrils that are straight, rigid rods on length scales below 100 nm, which is the range of the simulations. (Created with BioRender.com)

Understanding how this access is modified by experimental protocols or natural variations in fibril structure due to post translational modifications or unnatural modifications that are commonly introduced to facilitate the detection and monitoring of these proteins in cells and in vivo, is a prerequisite for a better understanding of fibril involvement in disease. Here we use molecular simulations to reveal how the accessibility of the surface of an amyloid fibril composed of the protein alpha synuclein is modified when the individual monomers are tagged by Green Fluorescent Protein (GFP).

Alpha synuclein (aSyn) is an intrinsically disordered protein (IDP) that is genetically and biochemically linked to Parkinson’s disease (PD) pathology and pathogenesis.[16, 17] aSyn fibrils are one of the main protein components of the pathological inclusions, called Lewy Bodies, found in the brain of PD and other neurodegenerative diseases that are collectively referred to as synucleinopathies, including Dementia with Lewy Bodies.[2] aSyn is a 140-residue sequence with disordered N and C terminal domains. Under pathological conditions it spontaneously assembles into rigid fibrils whose growth is enhanced by interactions of the monomeric protein with the growing ends or lateral surface of the fibrils.[18] Cryo-EM studies have shown that aSyn fibrils are composed of two protofilaments, and have diameters around 10 nm and lengths of 20-500 nm (see Figure 1).[19] The protofilaments contain a rigid core formed of the central part of the protein while the final ∼40 residues at the N and C termini extend in a disordered manner into the surrounding fluid. The C-terminal region is highly negatively charged, harbors most of the disease-associated post-translational modifications (PTM) of the protein and represents an exposed interactome hub. Increasing evidence suggests that many of these PTMs occur post fibrillization and contribute to the packing of fibrils into dense aggregates within Lewy Bodies.[4]

Experiments on fibril growth and interactions in cell models and *in vitro* frequently attach fluorescent labels to amyloid-forming proteins to make them visible in microscopy.[20] These labels include ligand-coated gold nanoparticles,[21] with diameters in the range 2.5 – 4 nm, and GFP,[22] with a linear dimension of 4 nm. GFP is typically covalently bound to fibril monomers by flexible peptide linkers containing 10-15 residues (equivalent to several nm),[23] usually to sequences that decorate the surfaces of the fibrils.[24] These sizes are comparable to the fibril diameters, which are in the range of 5-10 nm, allowing the morphology of the fibrils to be visualised. Figure 1 shows a cartoon that illustrates our hypothesis that the linked GFP tags partially occlude the surface of the fibrils and may interfere with biochemical reactions at its surface, including secondary nucleation and interactions with soluble proteins or other molecular species. The cryo-EM image in Figure 1 shows that the fibrils are typically rigid straight rods on length scales below ∼ 100 nm. We note here that the disordered termini of the aSyn protofilaments also protrude from the fibril surface and may also modify its accessibility, but this is not shown in Figure 1 for clarity and we do not address their effects in this work.

The process by which a diffusing molecular species interacts at a fibril surface consists of two sequential steps: 1) the molecules approach the fibril by diffusion; 2) closely-apposed molecules undergo conformational fluctuations leading to a complex, time-dependent interaction with the surface. The presence of the GFP tag interferes with the first process because it hinders diffusion to the fibril surface. Whether it affects the second process is a complex function of the diffusing particle’s size and conformational ensemble, and the GFP linker length. We do not attempt to address the full growth process here, but use coarse-grained molecular simulations to study the first stage in which untagged monomers diffuse to a fibril’s surface. Computer simulations provide a powerful tool to explore the kinetics of interactions of diffusing particles, oligomers, and amyloid fibrils.[25] However, many soluble disordered proteins have hydrodynamic radii of a few nanometers and require hundreds of nanoseconds to diffuse distances comparable to the diameter of aSyn fibrils.[26] Lipid vesicles and organelles are larger still, with diameters in the range of many 10s of nanometers. This makes the use of Atomistic Molecular dynamics computationally prohibitive, although it has been used to explore conformational fluctuations of single disordered proteins.[27] To retain near-molecular detail, we use the coarse-grained technique of dissipative particle dynamics[28-30] in which a single aSyn fibril is represented as a rigid cylinder to which GFP tags can be attached by flexible linkers (Figure 1). The linkers represent peptide chains with sizes in the range 2 – 4 nm (∼ 7 – 15 residues), but we do not assume a specific sequence beyond its flexibility, which typically requires glycine, proline or serine residues.[23] The diffusing monomers are modelled as spheres and ellipsoids with a hydrodynamic radius comparable to those of monomeric aSyn.[26] Although these proteins have a large conformational ensemble in solution,[31] their experimental diffusion is frequently modelled by spheres of an equivalent hydrodynamic radius.[26, 32] We do not attempt to capture atomistic details of these particles, which allows them to represent any diffusing object of equivalent hydrodynamic radius. These could be aSyn monomers or other proteins known to interact with its surface, small lipid micelles or membranous clusters.

Our results show that fluctuating GFP tags significantly reduce the residence time of diffusing particles at the fibril surface, and the magnitude of the effect has a complex dependence on the particle size and linker length. A non-monotonic variation in the residence time is found for particles whose hydrodynamic radius is 1-2 nm as the GFP linker length increases. Although the GFP tags hinder the approach of small particles to the fibril surface, once they have reached it they stay longer in its vicinity because the tags retard their diffusion away. Finally, we caution that experiments investigating the growth, interactome, and toxicity of GFP-labelled amyloid fibrils should be interpreted with care to eliminate the artifacts arising from the complex occlusion of the fibril’s surface by the tags.

## Results

### How do particle size and linker length influence fibril surface accessibility?

The physical dimensions of GFP tags are comparable to those of the hydrodynamic radius of monomeric intrinsically-disordered proteins such as aSyn. Our first aim is to explore how the presence of tags on a pre-formed fibril affects the ability of diffusing particles to interact with the fibril surface. Cryo-EM data from Guerrero-Ferreira et al.[19] is used to set the diameter of the fibril to 10 nm and the in-register monolayers have thickness 0.5 nm representing pairs of protofilaments. The fibril is circular because the atomic structure of its surface is not observable at the resolution of the coarse-grained simulations (∼ 1 nm). We use the explicit-solvent, coarse-grained simulation technique of dissipative particle dynamics (Materials and Methods section 1) to follow the diffusion of small nanoparticles around the model aSyn fibril and measure the time they spend near its surface. A single fibril is preassembled in the centre of the cuboidal simulation box with its long axis oriented along the Z axis of the box. A GFP tag may be attached via a short flexible linker to the C-terminus of each monomer in the fibril (**Figure 2)**. The remainder of the box is filled with solvent particles and a number of rigid diffusing particles as described in Section 2 of Materials and Methods. These particles represent untagged, monomeric aSyn proteins in dilute solution around the fibril. The hydrodynamic radius of aSyn is approximately 3 nm,[26] and this is taken as the upper limit of the particle diameter. We expect that representing an IDP by a nanoparticle with the same hydrodynamic radius is reasonable for studying the kinetics of their diffusive approach to a fibril in the presence of sterically-repelling bound GFP tags. Each simulation is run for 4 million time-steps, and the first 2 million steps are discarded to ensure that the system is in equilibrium before we take measurements.

**Figure 2.**
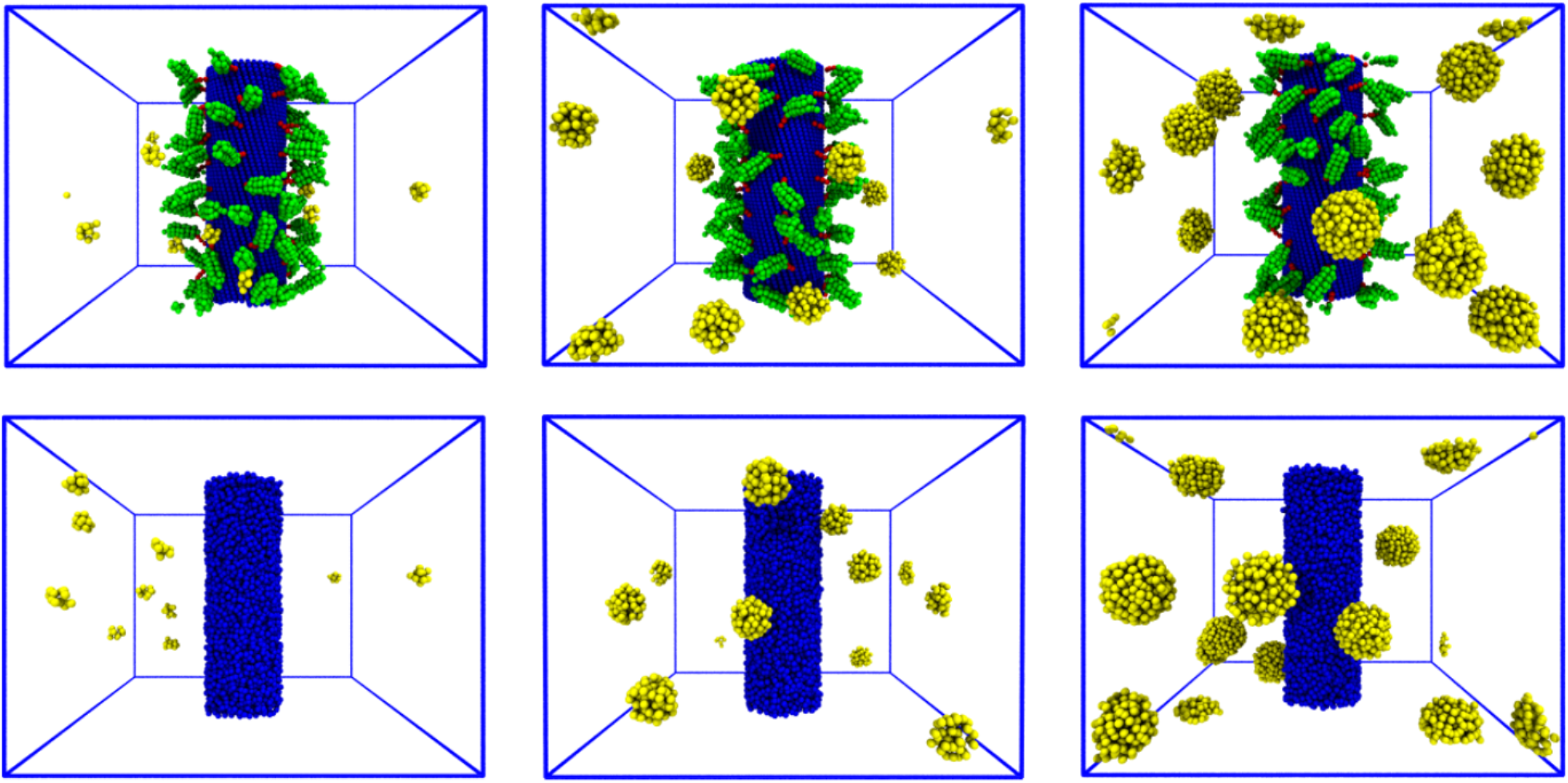
Snapshots from simulations of a 30 nm long fibril of diameter 10 nm in a box of size 40 × 40 × 30 nm^3^ with 10 spherical nanoparticles diffusing in the bulk solvent (solvent particles are invisible for clarity). The GFPs are modelled as rigid cylinders of size 3 × 4 nm attached to the fibril by 2 nm long flexible linkers. Top row shows the tagged fibril with spheres of radii 1 nm (**left**), 2 nm (**middle**), and 3 nm (**right**). The bottom row shows the bare fibril with the same number of nanoparticles that is used to define the baseline surface accessibility. The wide variety of conformations accessible to the GFP tags due to the flexible linkers is clear. Particles apparently cut by the simulation box boundary are connected via the periodic boundary conditions.

The bottom row of snapshots in Figure 2 show 10 spherical particles of radius 1 nm, 2 nm, and 3 nm (left to right) diffusing around the bare fibril. The upper row shows the same cases with the fibril decorated with rigid GFP molecules via 2 nm flexible linkers. Supplementary Figure S1 shows the equivalent case for 4 nm linkers. The GFP tag is represented as a rigid cylinder of dimensions 4 × 3 nm.[22] Thermal motion of the tags causes them to fluctuate over an area greater than their own dimensions depending on the linker length, an effect not apparent in the static snapshots in Figure 2 but visible in the Supplementary Movie SM1. Supplementary Figure S2 shows snapshots from the equivalent simulations for ellipsoidal nanoparticles of similar dimensions.

Previous computational studies have shown that globular domains, including GFP, linked to model Amyloid Beta monomers by short peptides interfere with their ability to oligomerise by creating a steric hindrance between monomers.[33] The linker used in this study was a flexible Ser-Pro-Ser chain. A minimal length of 7 residues (∼ 2 nm) was predicted to allow fibril elongation while ensuring the GFPs did not sterically intersect. In that work, the GFP molecules bound via a linker to the fibril monomers were stationary in space. However, the flexible peptide linkers used in experimental assays allow the GFPs to fluctuate around the fibril core.[23] In the case of aSyn, previous experimental studies have shown that the fusion of GFP to the C-terminus of aSyn with linkers ranging from 6-13 amino acids did not interfere with the ability of the protein to form fibrils *in vitro*.[34, 35] In this study, we have estimated the consequence of thermal fluctuations on the occluding power of the tags as follows.

The ease with which the particles can approach the fibril surface by diffusion is intuitively expected to be harder when the GFP tags are present compared to the bare fibril. We define a surface occlusion factor as the ratio of the amount of time the diffusing particles spend within a fixed distance of the fibril core with the GFP tags present to the time spent close to the bare fibril, normalised to remove the effects of the number of particles and simulation time. It represents the change in the equilibrium probability for the particles to be at the fibril surface. We quantify this measure by calculating the cumulative probability for the particles to be within a fixed distance of the fibril with and without the decorating GFPs (Materials and Methods section 3). The accumulated time spent by all the diffusing particles in concentric circular shells of equal radius centred on the fibril both with and without attached GFP tags is measured in independent simulations. These times are normalised by the total simulation time and number of particles, and the area of each shell. Figure 3 shows the probability histograms corresponding to particles of radius 1 and 2 nm shown out to 20 nm from the fibril centre, which is half the width of the simulation box (the full histograms are shown in Figure S2 of the Supplementary Material). The bottom row shows that the probability to find particles around the fibril is uniform beyond the minimum distance of the sum of the fibril and particle radii R_fibril_ + R_particle_. Results are not shown for particles of radius 3 nm because at a concentration that generates results of reasonable accuracy, the crowding of the particles in the simulation box modifies the probability distribution so that it cannot be compared with those of the smaller particles (see Supplementary Movie SM2). We expect that the displacement of the particles away from the fibril seen for 2 nm radius particles will be even stronger for larger particles.

**Figure 3.**
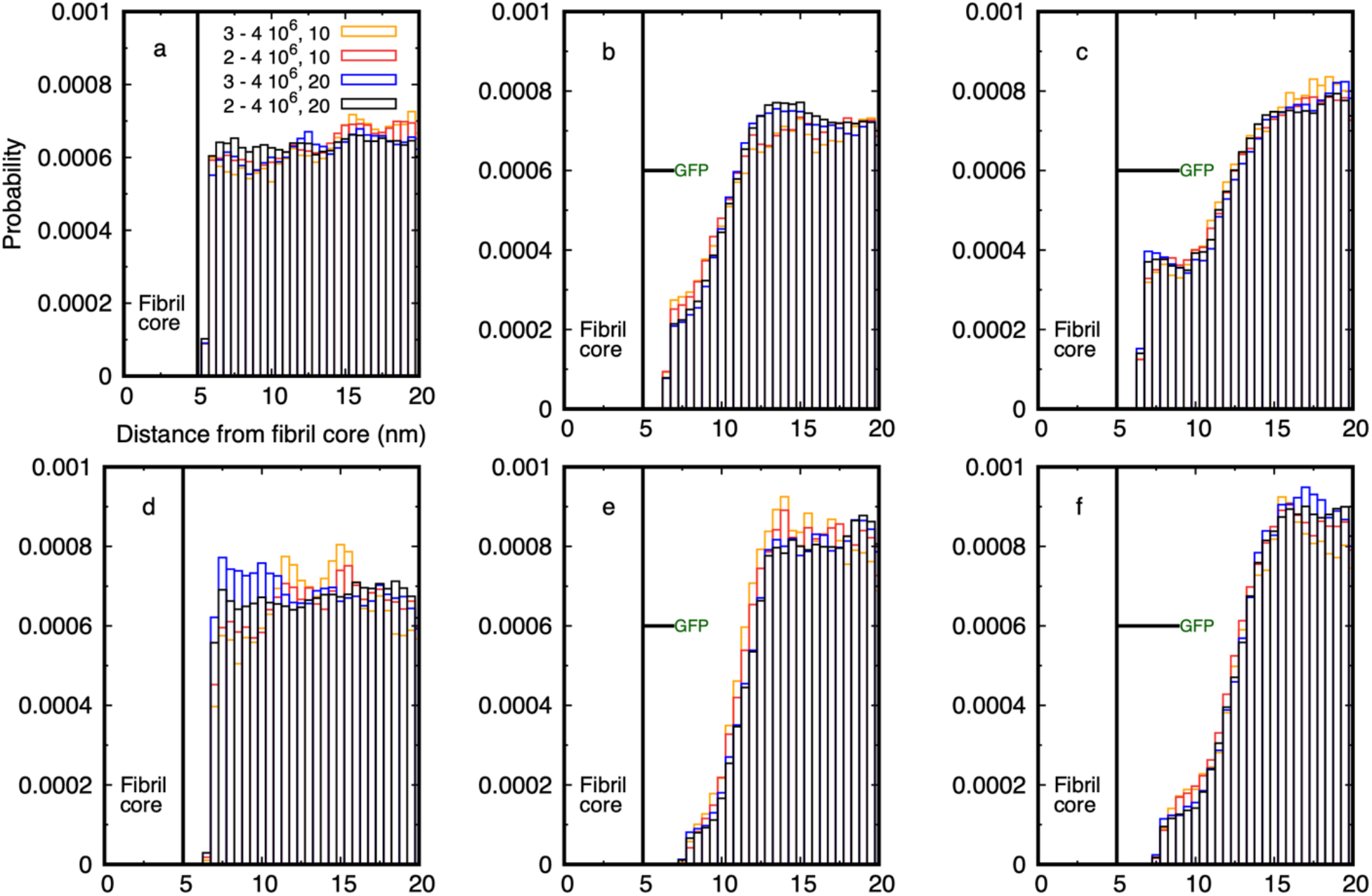
Histograms of the probability for diffusing spheres to have their centre of mass in cylindrical shells around the fibril of radius 5 nm. Four histograms are shown to illustrate the statistical accuracy of the distributions. The first two million time-steps of each simulation are discarded, and the distributions are sampled from 2 – 4 10^6^ and 3 – 4 10^6^ time-steps for 10 and 20 particles. The legend applies to all graphs. The left column is for a bare fibril and the GFP linker length in the middle and right columns is indicated by the horizontal line. Only one GFP is shown on a straight linker for clarity, whereas every monomer in the simulated fibril has a linked GFP and thermal motion causes the linkers to fluctuate in space. **(a)** The probability distribution for spheres of radius of 1 nm around a bare fibril is flat for all space outside the fibril (cp. left column of Figure 2). (**b**) The distribution for 1 nm radius spheres around a fibril decorated with GFP on a 2 nm linker shows a reduction in near-fibril probability as the fluctuating tags partially exclude the particles from the core. (**c**) The distribution for 1 nm radius spheres around a fibril decorated with GFP on a 4 nm linker shows the particles have an enhanced probability to be close to the fibril as the tags retard their diffusive escape. (**d**) The distribution for spheres of radius of 2 nm around a bare fibril is similar to (a) except it is displaced further out by the sphere radius (cp. middle column of Figure 2). (**e**) The distribution for 2 nm radius spheres around a fibril decorated with GFP on a 2 nm linker shows a larger reduction in near-fibril probability compared to the 1 nm spheres in (b). (**f**) No enhancement in the near-fibril probability is seen for 2 nm radius spheres when the GFP linker is 4 nm indicating that thermal fluctuations of the tags prevent the larger spheres from approaching the fibril surface.

As shown in Figure 3, the probability for a particle of radius 1 or 2 nm to be found between the fibril surface and the GFP tag is strongly reduced below that of the bare fibril. However, this drop is non-monotonic as the linker length is increased for the smaller particle. Although linkers of both lengths reduce the probability for the R = 1 nm particles below the bare fibril value, the longer linker results in a smaller reduction compared to the short linker, and there is a small peak at a separation of about 7 nm from the surface (compare middle and right histograms, upper row). There is no equivalent peak for the 2 nm radius particles, and the probability for particles to be nearer the surface than the GFP is still very small, although slightly larger for the 4 nm linker than the 2 nm one (compare middle and right histograms, lower row). Supplementary movie SM3 shows typical simulations of 20 spherical nanoparticles of radius 2 nm for the case of GFP attached by 2 nm and 4 nm linkers.

### Dependence of fibril surface accessibility on particle shape

The degree of occlusion of the fibril surface most likely depends on the shape as well as the size of the diffusing particles. This is seen in Figure 4 which shows the probability histograms for 10 and 20 spheres and ellipsoids with similar dimensions (Supplementary Movies SM4 and SM5). The distributions are similar for both types of particles when the fibril is bare and for the smaller particles with both linker lengths (upper row of Figure 4), although the large ellipsoids show an enhanced probability near the fibril surface as their smaller transverse dimension allows them to orient along the surface. Note that the smallest ellipsoids have dimensions 1.5 × 1 nm, which is quite similar to spheres of radius 1 nm. Given the resolution of the coarse-grained particles in the simulations, we are unable to construct ellipsoids of this size with a more precise shape.

**Figure 4.**
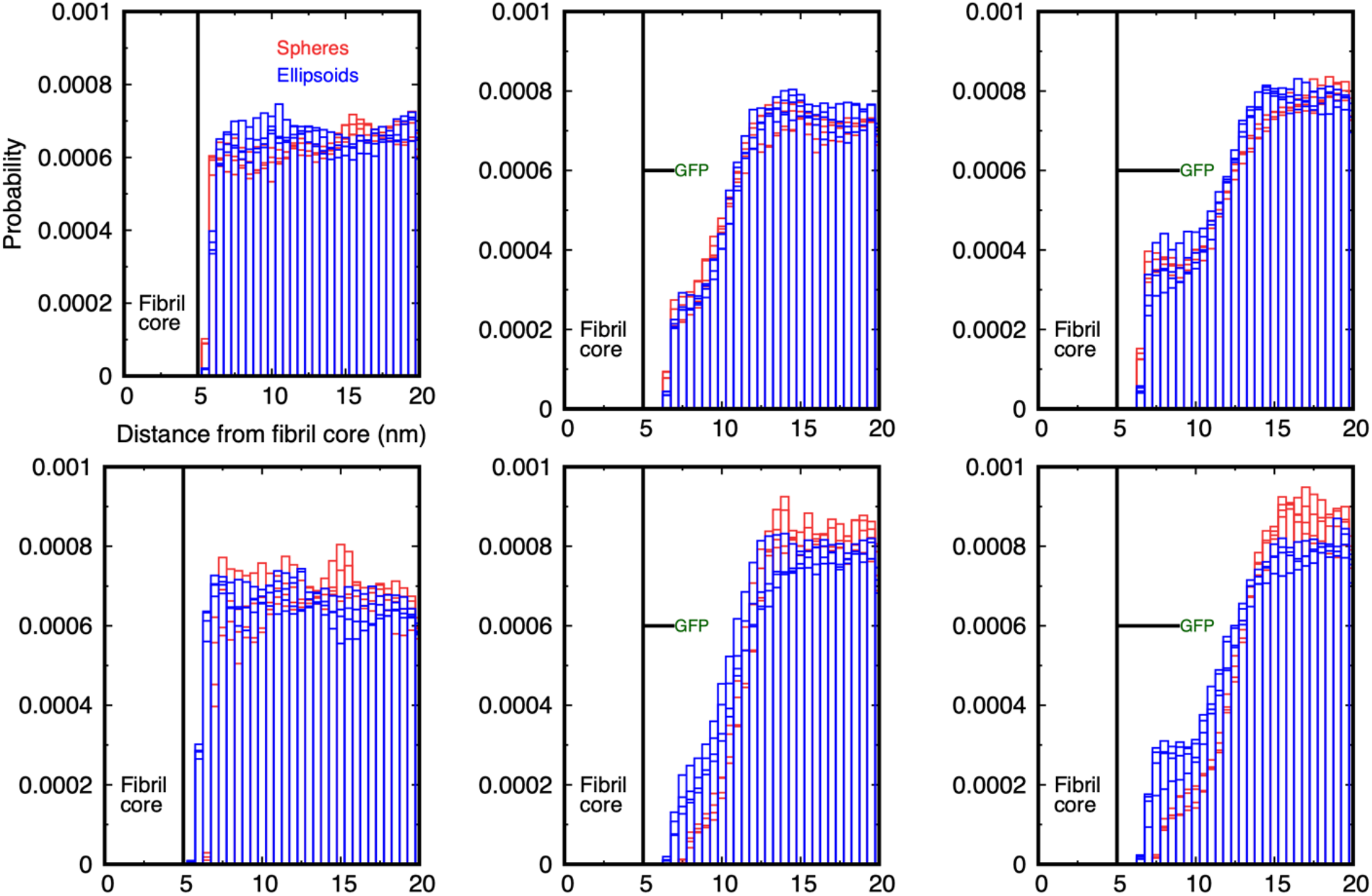
Comparison of the probability for spheres (red boxes) and ellipsoids (blue boxes) to have their centre of mass in cylindrical shells around the fibril. The first two million time-steps of each simulation are discarded, and the distributions are sampled from 2 – 4 10^6^ and 3 – 4 10^6^ time-steps for 10 and 20 particles. The legend applies to all graphs. **(Top row)** Probability distribution for spheres of radius 1 nm and ellipsoids with size 1.5 × 1 nm around the bare fibril (left), GFP-tagged fibril with 2 nm linker (middle), and GFP-tagged fibril with 4 nm linker (right). **(Bottom row)** Equivalent probability distributions for spheres of radius 2 nm and ellipsoids of dimension 2 × 1 nm. The distributions are similar in the bare fibril case, but the ellipsoids have an enhanced probability to be near the fibril surface compared to spheres of comparable size when the GFP is attached by the 2 and 4 nm linker, although the effect is smaller than for the smaller particles. Note that the larger ellipsoids are also able to approach closer to the fibril surface than the spheres because of their asymmetric shape.

The larger ellipsoids, of dimensions 2 × 1 nm, show an increased probability to be near the fibril for both linker lengths, the difference being larger for the longer linker (lower row of Figure 4). This indicates that the ellipsoids are able to diffuse to the surface more easily in the presence of the tags than spheres with equivalent dimensions. This result is expected geometrically as an ellipsoid whose semi-major axis is equal to the diameter of a sphere is smaller in the transverse dimensions, and can more easily diffuse between the tags. Molecular dynamics simulations predict that aSyn and other IDPs sample a wide range of conformations in solution,[36] which suggests that it may be inaccurate to use their equivalent hydrodynamic radius to characterise their diffusion in the presence of GFP-decorated fibrils.

We have shown that the presence of GFP tags attached to a fibril by flexible linkers of length 2 and 4 nm reduces the accessibility of its surface to diffusing particles in all cases studied compared to the bare fibril. The range of linker lengths and particle sizes examined is comparable to the hydrodynamic radius of monomeric aSyn and similar IDPs. We quantify the surface occlusion as follows. The particles are sterically unable to penetrate the fibril and, far from the surface, are relatively unaffected by the GFP tags. For a given linker length and particle radius, there is a range over which the probability histogram is modified by the GFP tags. We integrate the probability over this range for the decorated fibril and the bare fibril and use their ratio as a measure of the occluding effect of the tags. This measure depends on both the linker length and particle radius, but is normalised to be independent of the number of particles and total simulation time (see Section 3 of the Materials and Methods).

Table 1 shows the baseline probability for particles to be within a fixed distance of the fibril surface (column 3), and the ratio of this probability integrated over the same range for both linker lengths is shown in columns 4 and 5. For spheres and ellipsoids of dimensions 1 nm, the short linker reduces the surface accessibility to 30-40% of its bare value, and the longer linker to 40-50%. This drops to 10% and 15% respectively for particles of dimension 2 nm. The values for larger spheres are too small to be significant as they are effectively completely excluded from the fibril’s surface, but are shown for completeness. However, ellipsoids of semi-major axis 2 and 3 nm show a greater probability of being near the surface than the equivalent size spheres. The enhancement in the surface accessibility as a result of increasing the linker length is defined as the ratio of the two occlusion fractions, and is shown in the final column. It is clear that for spheres and ellipsoids, the enhancement can be large. The 1 nm spheres can spend more than 45% more time closer to the fibril surface when the linker is 4 nm compared to 2 nm, and ellipsoids of size 2 and 3 nm can spend up to 80% more time at the surface, albeit from a lower baseline. Note that although the final column of Table 1 shows that spheres of radius 2 nm have a greater enhancement at the surface for the longer linker than the 1 nm spheres, Figure 3 and the third column of Table 1 show that they actually spend much less time there than the 1 nm spheres, and the apparent increase is from a lower baseline value.

**Table 1.**
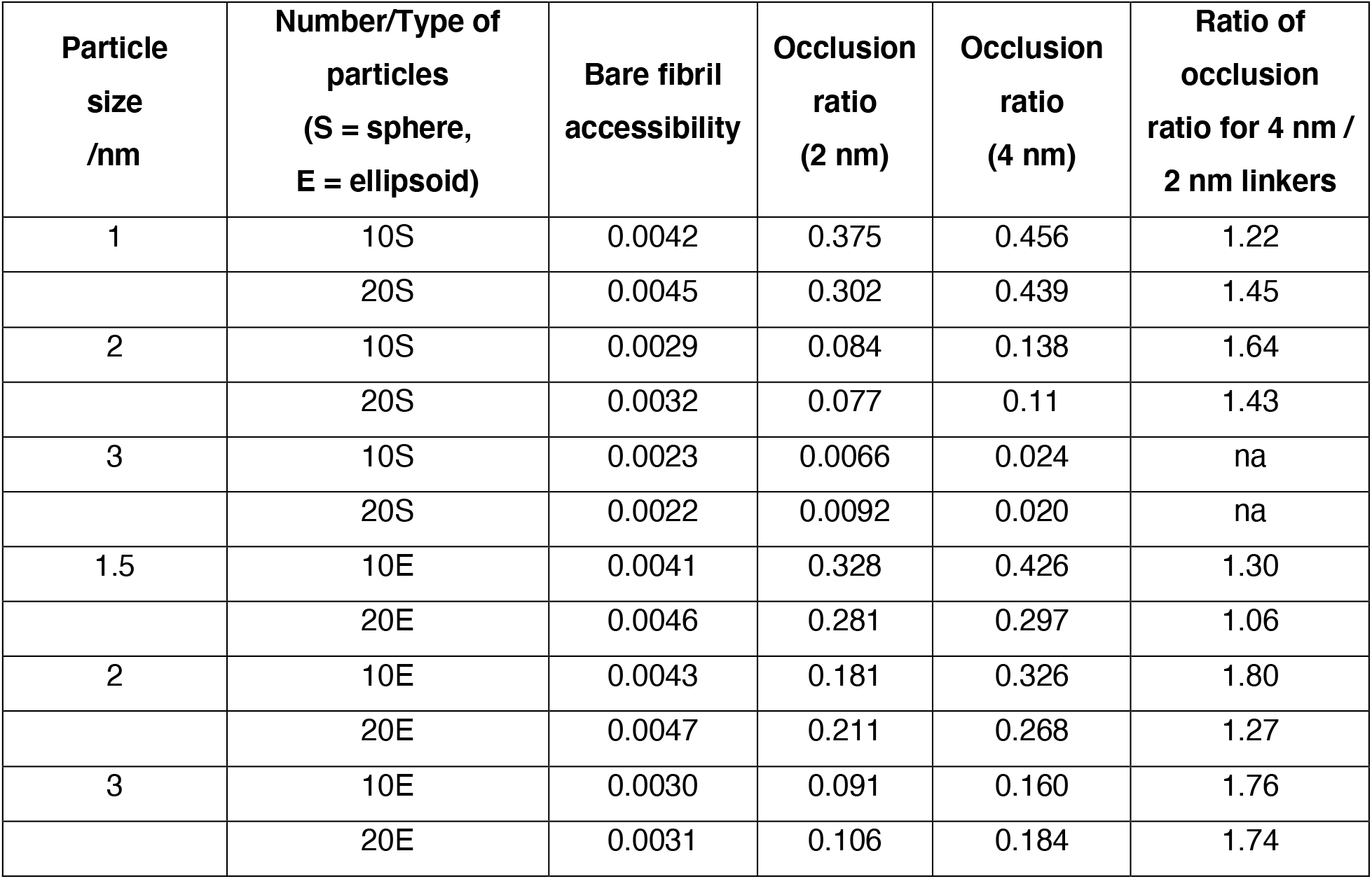
Effect of the linker length on the fibril surface accessibility for spherical and ellipsoidal particles of different sizes. The first column gives the radius of the spherical particles and the semi-major axis of ellipsoidal particles. The fibril diameter is 10 nm in all cases. The bare fibril accessibility is the integral of the histogram for the particles to be within 4 nm of the fibril surface (9 nm from its axis, cf. Fig. 3). The occlusion ratio is the fraction of the bare fibril accessibility remaining when the linker/GFP combination is present: a value of 1 means there is no occlusion while a value of 0 means access to the surface is entirely blocked. These are values taken from independent simulations of each type. The final column shows the occlusion ratio for the 4 nm linker case divided by that for the 2 nm case, and quantifies the magnitude of the enhancement due to the larger space behind the GFP on the long linker. The occlusion ratios are calculated by integrating the radial distribution function from the fibril surface out to a given distance as described in the Supplementary Material. na = not applicable as the values are too small to be determined accurately.

It might be expected that the occluding effect of the tags would decrease with increasing linker length because of the greater free space around the fibril by which diffusing particles can approach its surface. We have tested this hypothesis by performing simulations in which the GFP tags are connected by linkers that are 10 nm long in a larger simulation box (50 × 50 × 30 nm^3^). Because these simulations are computationally expensive, we present only the histograms of the probability distributions of particles around the fibril. Figure 5 shows that the fibril surface is still significantly occluded for particles of radius 1, 2, and 4 nm even when the linker is 10 nm long. The figure shows that although the 1 nm radius particles are able to penetrate to the fibril, those with a radius of 4 nm are unable to diffuse closer than 5 nm of its surface. Supplementary Movie SM6 shows the three cases of the particles of different radii diffusing around the tagged fibril.

**Figure 5.**
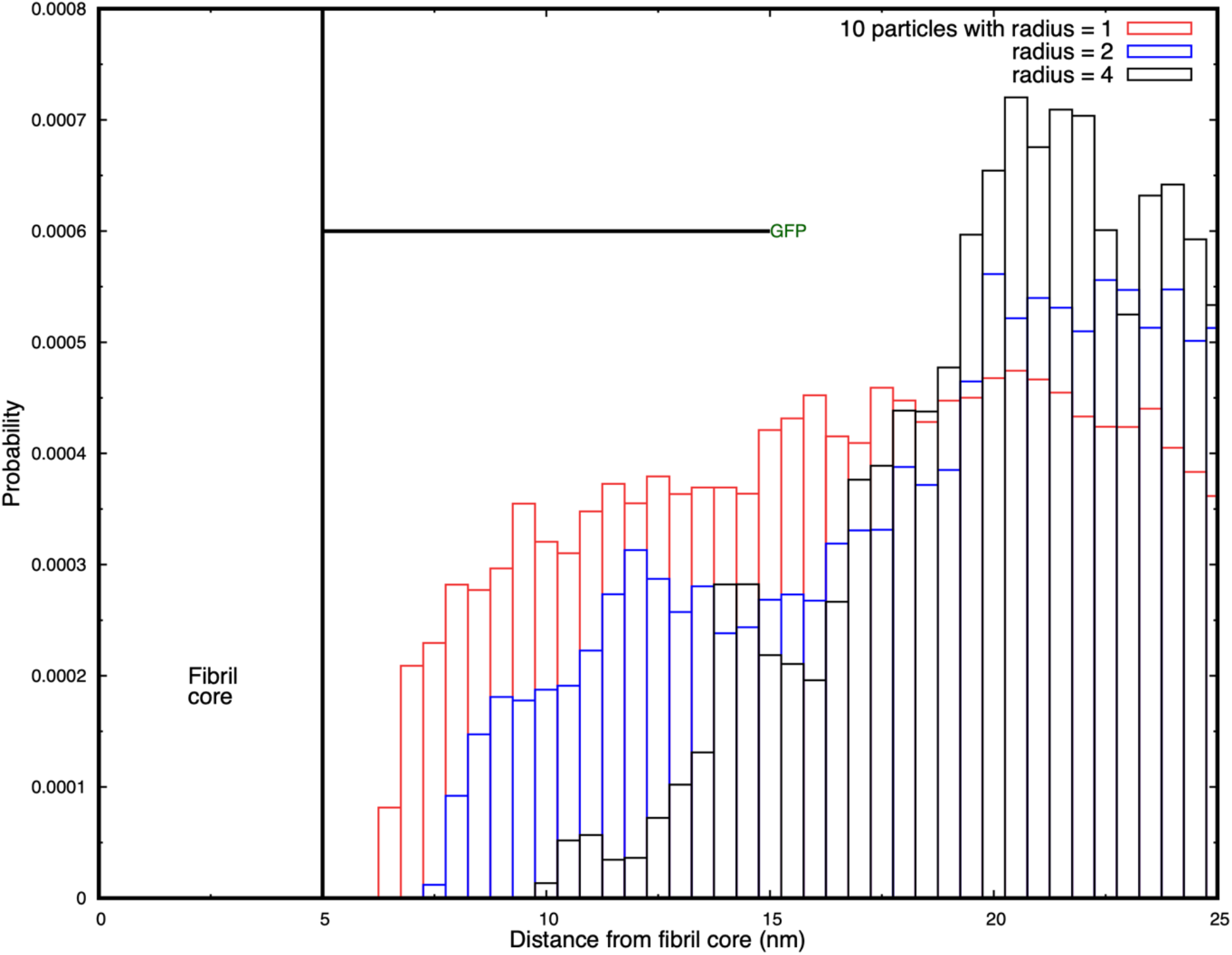
Histograms of the probability for diffusing spheres to have their centre of mass in cylindrical shells around the fibril. Results are shown for 10 particles of radius 1, 2, and 4 nm, and the occlusion increases with increasing particle size. The first two million time-steps of each simulation are discarded, and the distributions are sampled from 2 – 4 10^6^ and 3 – 4 10^6^ time-steps.

## Discussion

We have used coarse-grained simulations to explore how fluorescent protein tags bound to a model aSyn fibril by flexible linkers occlude its surface to particles approaching by diffusion. Soluble proteins such as monomeric aSyn are treated as rigid spheres or ellipsoids that diffuse with the same hydrodynamic radius as that obtained experimentally. The accessibility of the fibril surface is always reduced by the presence of bound fluorescent tags compared to the bare fibril, as expected intuitively, and the effect is generally stronger as the particle size and linker length increase within the range of 2 – 4 nm. However, the surface occlusion is a non-monotonic function of the GFP linker length and particle size, and the residence time of small particles at the surface is counter-intuitively *increased* by the presence of GFP tags when the particles are smaller than the linker length. The linker lengths used to attach fluorescent groups to monomers in amyloid studies are often 10 – 14 residues, which corresponds to 3-4 nm (taking the average length of a residue to be 0.3 nm) and this is the range we have studied here.

Surface occlusion occurs because thermal fluctuations of the GFP tags around the fibril surface create a steric barrier to the approach of diffusing particles. We hypothesize that the increase in residence time for particles smaller than the linker length occurs because although the GFP tags interfere with the approach of the particles to the fibril, they also transiently hinder their diffusive escape. This results in a significant enhancement in their residence probability at the surface. We find that this is not an insignificant effect: spheres of radius 1 nm spend ∼50% more time at the surface when the GFP is attached by a 4 nm linker than for a 2 nm linker. Ellipsoidal particles show a similar pattern of enhanced residence at the surface. Compared to previous work in the literature in which the steric effects of static tags were explored,[33] we find that thermal fluctuations of the tags gives rise to more complex occlusion of the fibril surface.

The question arises how relevant it is to approximate the steric interactions of a conformationally-fluctuating, soluble protein by those of a rigid sphere or ellipse? Although intrinsically-disordered proteins are not rigid, as are folded proteins, their hydrodynamic radius is often used to calculate their diffusion. Marsh and Forman-Kay[32] and Tomasso et al.[26] have tabulated how the hydrodynamic radius of an IDP scales with the number of residues. An aSyn monomer has 140 residues, and is predicted by Tomasso et al. to have a hydrodynamic radius of 2.7 nm. This may be compared to experimental values of 3.17 nm (pulse-field gradient NMR)[37] and 3.27 nm (FCS).[38] Experimental values depend on pH, the molecule becoming more compact at lower pH. The relation between the hydrodynamic diffusion of an IDP and its molecular size is therefore also of experimental interest. The steric interactions of a fluctuating polymer arise from direct contact of its monomers whose average spatial distribution is described by its radius of gyration. Dunweg et al. used Monte Carlo simulations to show that the radius of gyration of a self-avoiding polymer is 60% larger than its hydrodynamic radius.[39] This is in contrast to a uniform sphere, for which the radius of gyration and hydrodynamic radius are related by the familiar formula 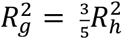. This implies that the steric interactions of fluctuating disordered proteins are *larger* than those of a uniform sphere of the same hydrodynamic radius. Steric interactions between soluble monomeric IDPs and fibrils in experiments are likely to be greater than those present in our simulations, which represents a lower limit to the occluding effect. Other typical fluorescent dye molecules such as rhodamines, oxazines, and fluorescein have dimensions around 0.7 – 1 nm, but can be larger when attached to proteins. These are smaller than GFP and are expected to produce a smaller occluding effect.

Our results also allow us to predict how the occlusion affects the rates of reactions of soluble proteins at the fibril surface. These rates are commonly used in constructing models of fibril nucleation and growth.[13] The rate constant of a unary reaction in which a protein interacts with the surface is predicted to drop to ∼45% of its well-mixed value for an untagged fibril for a protein with a 1 nm hydrodynamic radius, and to 10% for 2 nm radius. A binary reaction requires two proteins to meet at the surface. But we have only the single-particle probability to be within a 2 nm radius of the fibril’s surface. If we assume the probability distribution has translational symmetry along the fibril and circular symmetry around the fibril, and the diffusion of two particles to the surface are independent events, then the probability of two soluble proteins being within any small volume (of the size of the proteins) is the square of the single-particle probability scaled by the ratio of the small volume to the volume of the circular shell around the fibril. As the latter ratio is the same for the bare fibril as the tagged one, the reaction rate constant is just reduced by the square of the single-particle probability. Therefore, the rate constant will be reduced to 0.45^2^ ∼ 0.2 of its untagged value for 1nm radius proteins, and to 0.1^2^ ∼ 0.01 for 2 nm radius proteins. These are large reductions and should be apparent in experiments even given the statistical errors in the simulations. In our work, we have assumed that every monomer in the fibril is tagged by a GFP moiety. If there are gaps because of fewer tagged monomers, this will increase the surface accessibility and modify these predictions.

### Implications for the mechanisms of pathological aggregate formation and toxicity

Our modeling and experimental observations suggest that the presence of GFP on the surface of the fibrils could significantly modify their surface properties, the potential for post-translational modifications, and interactions with other proteins and organelles. Recent studies from our group and others have shown that these processes play a central role in the biogenesis of pathological inclusion formation.[3, 4, 6, 12, 40] Below, we reflect on the implications of our findings on the mechanisms of amyloid formation and toxicity based on the experimental data available in the literature today. We attempt to explain how GFP could influence the properties of amyloid aggregates and our ability to model critical processes linked to the formation and maturation of pathological inclusions associated with PD and other neurodegenerative diseases.

### The fusion of GFP influences the biophysical properties of amyloid fibrils

Fibril growth is a complex process, and GFP tags may interfere with the primary elongation mechanism or secondary nucleation at the surface, or both. It is known that fluorescent tags modify the size distribution of oligomers of the Alzheimer Aβ peptide,[20] and may also modify its interactome by creating a steric hindrance to the approach of diffusing molecular species, and the lateral association between decorated fibrils, which is important for their pathological inclusion formation and maturation.[41]

Several amyloid proteins have been expressed and purified as GFP fusion proteins, including amyloid-beta (Aβ, 4 kDa),[42] alpha synuclein (aSyn, 14 kDa),[34, 35] Tau,[43] Tau fragments and mutant Huntingtin fragments (Htt, exon1 11 KDa).[6, 44, 45] Despite the fact that GFP is much bigger in size (27 kDa) compared to most of these proteins, it did not interfere with their ability to form fibrils *in vitro*,[43] or in cells,[4, 6, 45] except for Aβ, where the addition of GFP resulted in complete inhibition of Aβ fibril formation *in vitro*.[42] These observations have led to the use of GFP-fusion proteins to investigate many aspects of the kinetics and mechanisms of amyloid formation in cellular assays and animal models. The assumption in many of these studies is that GFP does not alter the biophysical properties of the final fibrils or their ability to evolve and mature to the final pathological inclusions found in the brain, amyloid plaque (Aβ), Lewy bodies (aSyn), or neurofibrillary tangles (Tau). However, recent biophysical studies showed that the fusion of GFP to full length Tau (Tau_FL-GFP) or the short repeat domain Tau containing a pro-aggregating mutation (Tau_RDΔK-GFP), separated with a 13-14 amino acids linker, significantly alters the β-strands packing within the fibrils.[43] In addition, atomic force microscopy (AFM) showed that the fibrils formed by Tau_FL-GFP were wider and characterized by the presence of an additional halo of height corresponding to the size of GFP (≈ 3 nm). Similar observations were made for mutant Httex1 fused to GFP (Httex1-GFP).[44] The increase of fibril width by 3 nm is also consistent with the size of the GFP protein. Similarly, the generation of aSyn-GFP fibrils *in vitro* resulted in a significantly delayed aggregation kinetics and the formation of wider fibrils.[46] In the study by Afitska *et al*.,[47] the authors even reported that the fusion of GFP completely inhibited the primary nucleation of aSyn. In all cases, the resulting fibrils do not share the morphological properties of the fibrils found in AD, PD or HD brains. These observations demonstrate that the GFP subunits decorate the surface of the fibrils, change the surface properties of the fibrils and limit access to their core structure, thus altering their interactome or ability to catalyse surface-mediated secondary nucleation events. We summarized in Figure 6 the biophysical and cellular influence of GFP on amyloid fibrilization and inclusion formation *in vitro* and in cells.

**Figure 6.**
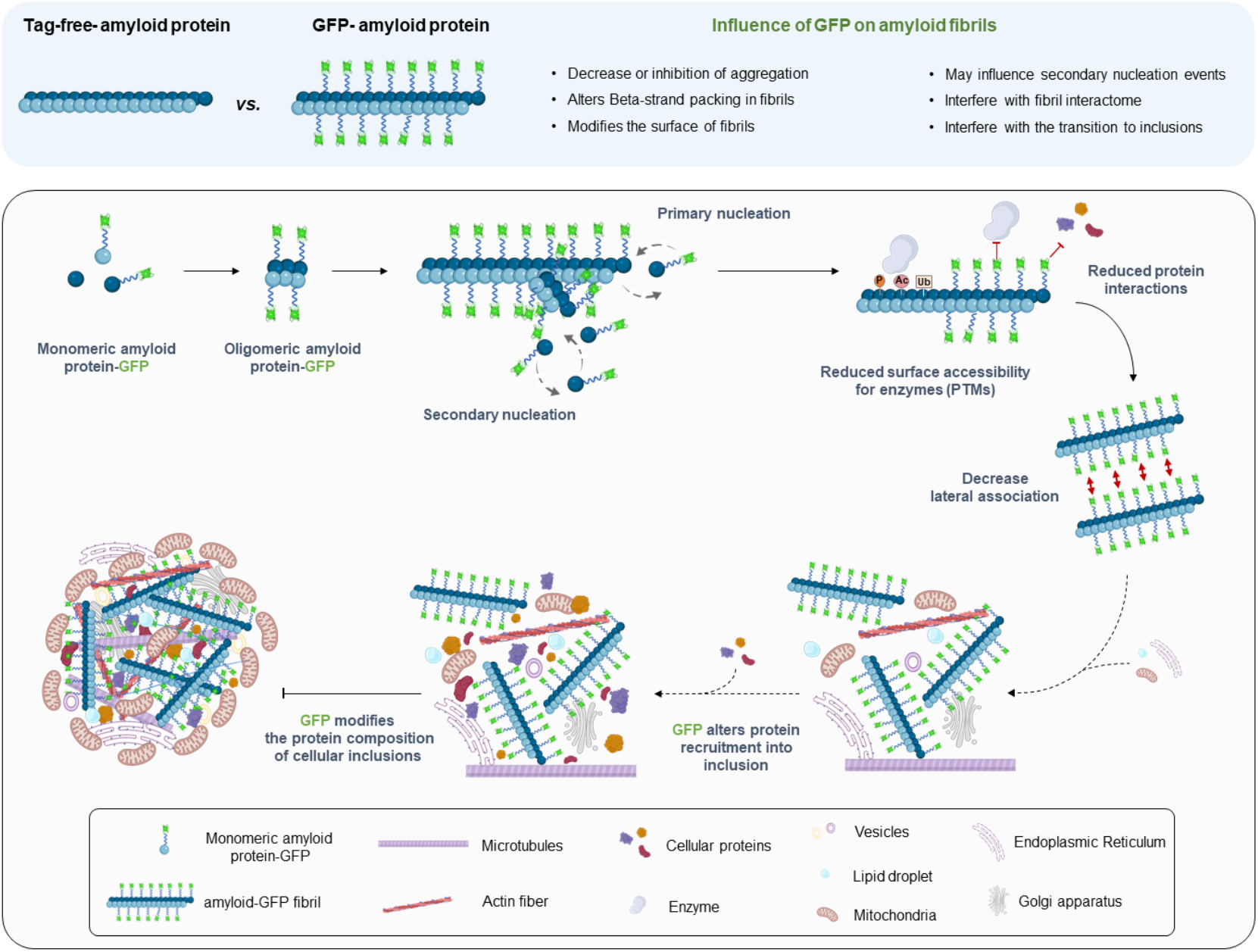
Schematic depiction illustrating the effects of GFP on the various stages of protein aggregation and inclusion formation based on published studies and predictions from our work. The depicted mechanisms illustrate the various stages associated with the mechanisms of aSyn oligomerization, fibrilization and LB formation.[3, 4]. (Created with BioRender.com.)

Many proteins are known to interact with aSyn fibrils[15] and their ability to access the fibril surface could influence fibril growth, post-translational modification, morphology and interactome, all of which influence their toxicity and ability to transition or mature into the pathological aggregates found in PD brains. Our results predict that the fluorescent tags act as a molecular “sieve” that differentially restricts access to the fibril surface for proteins of different sizes. We are not aware of any experimental data that compares the protein size-dependent interactome for aSyn fibrils with and without GFP proteins. However, we have recently determined the enrichment of soluble proteins in cytoplasmic inclusions in cells overexpressing mutant forms of exon 1 of the huntingtin (Htt) protein, with and without GFP fusion to their C-terminal domain. Our correlative light electron microscopy studies confirmed that Htt fibrils are the primary component of these inclusions, confirming that the presence of GFP does not interfere with the ability of mutant Htt to form fibrils.[48] Despite this, a careful comparison of the composition of Htt inclusions and their associated toxicity revealed that the presence of GFP strongly influenced the ultrastructure, proteome and lipid composition of the inclusions and their toxicity.

Given the availability of the proteome data for the Httex1 72Q and Httex1 72Q-GFP cytoplasmic inclusions from this study (https://www.ebi.ac.uk/pride/archive/projects/PXD021742), we sought to determine if the presence of GFP influences the size distribution of the proteins that are recruited into Htt inclusions in cells. Figure 7 shows the molecular weight distribution of proteins that are enriched in Htt inclusions in HEK cells. The right bar shows the composition in inclusions formed of bare Htt fibrils with a polyQ length of 72, which is above the value at which pathological fibrils form. The left bar shows the composition when the Htt fibrils are decorated with GFP tags at their C-terminal end. It is clear that while low molecular weight proteins are present in both cases, no proteins with masses above ∼ 150 kDa are found in the inclusions where GFP is present. Although the experimental situation is more complex than the simulations, it is consistent with the prediction of our simulations that GFP tags on flexible linkers preferentially exclude higher molecular weight proteins from the fibril surface.

**Figure 7.**
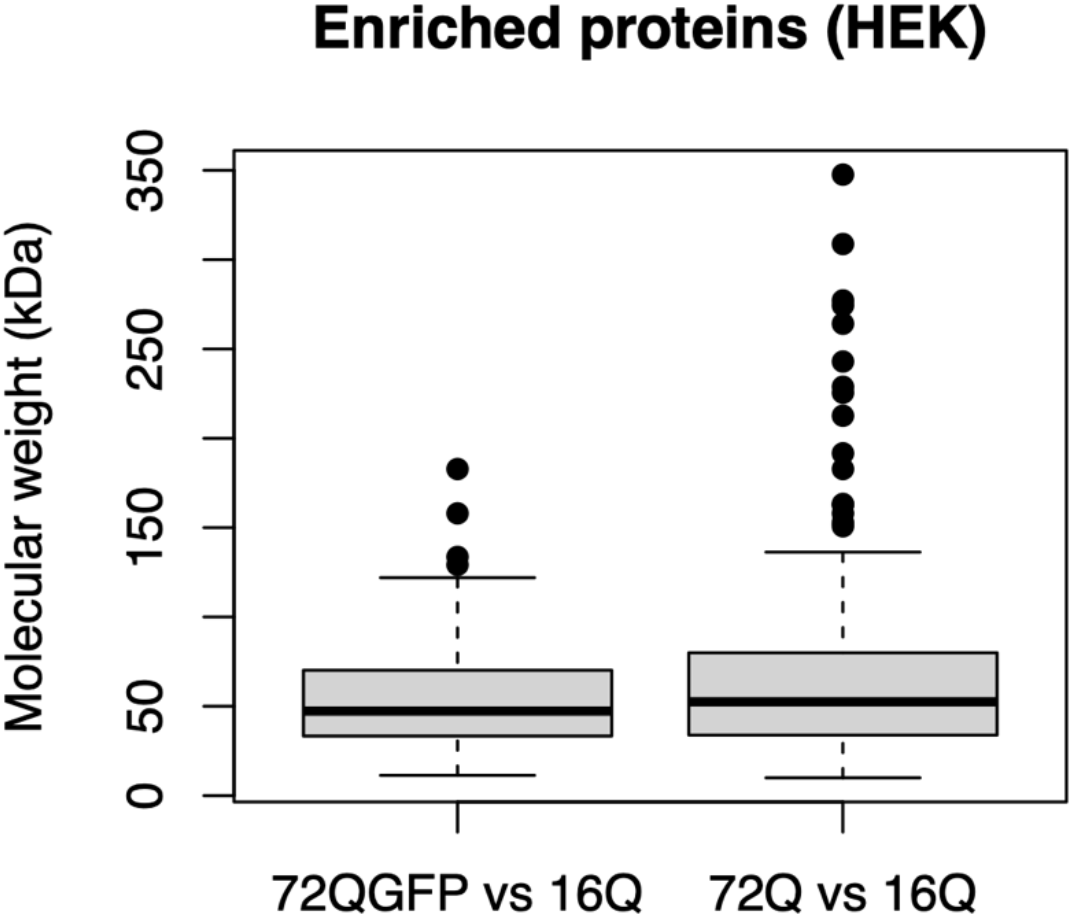
Comparison of the protein composition of cytoplasmic Htt inclusions in an experimental model of Huntington’s disease Htt aggregate inclusion formation from Ref. 6. HEK cells with inclusions formed after seeding with fibrils of Htt with a poly-Q repeat length of 72Q (as compared to a baseline of 16Q) decorated with GFP at the C terminus (left) and for unlabelled Htt (right). Inclusions containing label-free Htt fibrils are enriched in a wide range of proteins including many with high molecular weights. GFP-tagged Htt fibrils have a similar distribution of low molecular weight proteins (∼ 50 kDa), but there are no proteins with molecular weights above 160 kDa present in contrast to the label-free inclusions. The difference in the means between the two distributions is statistically significant (with a t-test for difference of means, p < 0.05) but what is most striking is the absence of proteins with high molecular weights in the GFP-tagged inclusions.

Finally, the presence of GFP on the surfaces of fibrils seems to also influence their self-association and packing during the formation and maturation of inclusions. Bäuerlein *et al*. reported that the fusion of GFP to the C-terminal part of Httex1 resulted in a 50% reduction in fibril density inside cytoplasmic inclusion formed in primary neurons and a 25% increase in fibril stiffness due to the GFP decoration along fibrils.[45] A review of the aSyn seeding models revealed that seeding mediated aggregation in cells or transgenic mice overexpressing aSyn-GFP resulted in the formation of filamentous aggregate but not LB-like inclusions.[34, 35] In Schaser *et al*.,[49] A53T mutant aSyn-GFP mice were injected with PFFs which resulted in the formation of pS129, and ubiquitin positive filaments but did not form round LB-like inclusions. Similarly, Trinkaus and colleagues performed cryo-electron tomography (cryo-ET) from neurons expressing A53T aSyn-GFP and treated with recombinant or brain derived PFFs.[50] In both cases, the detected aSyn neuronal aggregates were predominantly composed of aSyn fibrils in the middle of cellular organelles and membranes, but they did not observe Lewy-body-like spherical inclusions similar to those formed by endogenous untagged aSyn. These observations suggest that the presence of GFP interferes with aSyn fibril lateral association and interactions with cytoplasmic proteins and organelles, processes that are tightly linked to the formation of LB-like inclusions. Although aSyn-GFP can form fibrils, whether these fibrils can seed aSyn aggregation as native (tag-free) aSyn PFFs has not been investigated.[4, 51] Altogether, these observations and our data demonstrate that in addition to altering the kinetics of fibrilization of amyloid proteins and the biophysical properties of amyloid fibrils, the presence of large tags will also change the final structure and composition of protein aggregates and inclusions in cells.

### Implications for drug discovery and identification of modifiers of amyloid formation and clearance

High throughput screening of small drug molecules has uncovered compounds that interfere with the initial formation of aSyn fibrils or secondary nucleation events at the surface of existing fibrils.[52] The molecule ZPD-2 inhibits the initial seeding of new filaments and was found to be most active when added early in the aggregation reaction.[53] SynuClean-D, by contrast, is a small molecule that inhibits aSyn fibril aggregation and acts to disaggregate mature fibrils in human cell and *C. elegans* model systems, but does not strongly interact with monomeric aSyn. It was predicted by computational analysis to bind to small cavities in the fibril surface supporting the importance of the surface.[54]

Similarly, many of the therapeutic antibodies designed to target pathological aggregates or facilitate their clearance are designed to bind to sequences that decorate the fibril surfaces. Therefore, it is important that such therapeutic agents are validated in models expressing native protein sequences. The variation in the degree of occlusion with linker length and particle sizes suggests that experiments exploring the interactome of GFP tagged-amyloid fibrils with proteins or drugs, and the lateral association of multiple fibrils, should take account of the non-monotonic steric effects of tag and linker when comparing to label-free experiments. Additionally, we propose that *in vitro* experiments that measure reaction rates at fibril surfaces should be corrected for the effects of GFP tags when used to determine rate constants for theoretical modelling.[13] Accurately understanding fibril surface occlusion is therefore important for interpreting experimental data, building kinetic models of fibril elongation, and drug discovery studies.

Collectively, our results indicate that experiments that use tagged and untagged monomers to study the growth and interactome of fibrils should be compared with caution, and the confounding effects of the tags are more complex than a simple reduction in surface accessibility. The prevalence of fluorescent tags in amyloid fibril growth experiments suggests that this has implications beyond the specific alpha synuclein fibrils we model here. Finally, given the increasing use of cellular assays and biosensors based on the expression of the amyloid protein to fluorescent proteins in drug discovery, it is essential first to determine which aspects of the pathological protein aggregation process of interest are recapitulated in these assays.[55-61] This requires detailed characterization of the aggregates and inclusions formed at the ultrastructural and biochemical levels. Furthermore, compounds and antibodies identified using these assays should always be validated in cellular and animal models expressing untagged native proteins.

## Materials and Methods

### 1) Dissipative Particle Dynamics simulation technique

We use the Dissipative Particle Dynamics technique (DPD) to study the diffusive approach of rigid nanoparticles to a stationary model aSyn fibril. The source code for the DPD simulations carried out in this work is available on GitHub: https://github.com/Osprey-DPD/osprey-dpd. DPD is a coarse-grained, explicit-solvent molecular simulation technique designed to study the hydrodynamic behaviour of complex fluids,[28-30] and soft materials.[62-64] Its advantage over both atomistic and coarse-grained molecular dynamics are its speed of execution and retention of the correct hydrodynamic behaviour of the solvent. The speedup is obtained by grouping several atoms or atomic groups into *beads* that interact via soft forces. This allows a larger integration step size to be used in the equations of motion.

Atoms and molecular groups are represented by beads that interact via three non-bonded interactions that are soft, short-ranged (vanish beyond a fixed length-scale r_0_) and pairwise additive, conserving linear momentum. One force is conservative and gives each bead an identity such as hydrophilic or hydrophobic. Its magnitude is set by the parameter *a*_*ij*_, which is the maximum force between beads of type i and j. The other two forces are a dissipative and a random force that together provide a thermostat that maintains a constant system temperature. The magnitude of the dissipative force is set by the parameter *γ*_*ij*_. The masses of all beads are equal and set to unity. Molecules in DPD are constructed by connecting beads by Hookean springs defined by a spring constant *k*_2_ and unstretched length *l*_0_that may depend on the bead types. A bending stiffness potential may be associated with adjacent bonds in a molecule that has the form *k*_3_ (1 − *cos*(*φ*− *φ*_0_)1 where *k*_3_ is the bending constant (in units of k_B_T) and *Φ*_0_ is the preferred angle, which is zero if the bonds prefer to align parallel.

The aSyn fibril is composed of a central core made up of beads of type C that are constrained to be stationary; the linker is a short, linear chain of beads L, and the GFP moiety is a rigid cylindrical structure made of beads G. The solvent particles are represented by a single bead W that represents several water molecules. All the interaction parameters for the bead types and bonds are specified in Table 2.

**Table 2.**
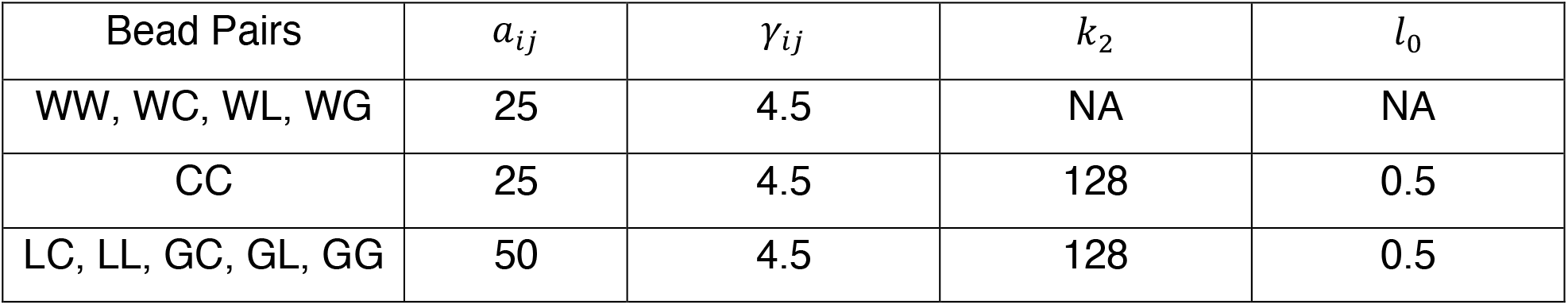
Bead-bead conservative force parameters *a*_*ij*_ (in units of *k*_*B*_*T*/*r*_0_) and dissipative force parameters *γ*_*ij*_ (in units of 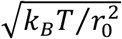) for all bead pairs, and Hookean bond potential parameters (in units of 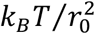 and *r*_0_respectively). The water beads have the same conservative interaction with all bead types. NA = not applicable. The bending stiffness parameters for the linker (LLL) and GFP beads (GGG) are *k*_3_ = 20 *k*_B_*T* and *φ*_0_= 0. Further details of the simulation parameters are given in the literature.[62, 64]

| Bead Pairs | $a_{ij}$ | $\gamma_{ij}$ | $k_2$ | $l_0$ |
| --- | --- | --- | --- | --- |
| WW, WC, WL, WG | 25 | 4.5 | NA | NA |
| CC | 25 | 4.5 | 128 | 0.5 |
| LC, LL, GC, GL, GG | 50 | 4.5 | 128 | 0.5 |

### 2) Constructing the fibril and nanoparticles

The simulation length scale is set by the experimental value for the aSyn fibril diameter, which we take as 10 nm and the monomer thickness 0.5 nm to correspond with cryo-EM experiments of aSyn fibrils.[19] The smallest nanoparticle has a 1 nm radius, and we set the range of the DPD non-bonded forces to this value, *r*_0_= 1 *nm*. A model aSyn fibril is preassembled in the simulation box from circular monomers that represent the paired aSyn protofilaments. The monomers are bound together with strong Hookean springs to give the fibril a high rigidity. Although aSyn fibrils have distinct polymorphs, including twisted structures,[65, 66] these sub-nanometer details are not visible in our coarse-grained simulations. We expect that the diffusive approach of a protein to the fibril surface is not greatly affected by atomic details of the surface until it approaches closer than one nanometer, which is below the accessible length scale here. We also ignore the disordered parts of the aSyn termini that protrude from the fibril’s surface.

The simulation box is 40 × 40 × 30 nm^3^ and the fibril is 30 nm in length oriented along the Z axis. This is a compromise between a sufficiently long fibril to minimise the effects of the boundary conditions at its ends and a reasonable computational cost of the simulations. The lateral dimensions of the simulation box are four times the fibril diameter also to minimise the effects of the system size on the results.

Because we are interested in equilibrium properties, we do not attempt to fix the simulation time scale precisely. But an approximate value can be obtained as follows. Stokes law predicts that the diffusion constant for a rigid particle undergoing Brownian motion in a medium is *D* = *k*_*B*_*T*/6πη*a* where *k*_*B*_ is Boltzmann’s constant, *T* is the temperature, *η* is the medium viscosity (0.001 Pa.sec for water), and *a* is the hydrodynamic radius of the particle. By comparing this with the dimensionless quantity 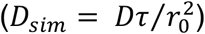 measured in the simulations, the DPD time-scale τ = 0.3 ns. Each simulation is run for 4 10^6^ steps with an integration step-size of 0.025 τ, which corresponds to 30 μsec, and requires 12 cpu-days on a single core of an AMD Ryzen Threadripper 3970X processor. The first 2 million steps are discarded to ensure the system is in equilibrium before we measure the observables.

When simulating the decorated fibril, each fibril monomer has a single GFP attached to it by a flexible linker. The GFP is represented as a rigid cylinder of diameter 3 nm and length 4 nm.[22] Successive GFP tags are rotated by 51 degrees in a spiral around the core.[67] The tags are sterically excluded from the fibril and each other. The flexible linker allows them to fluctuate in position in response to thermal noise subject to not intersecting other rigid objects in the simulation.

Finally, a number of nanoparticles are created in the solvent region of the box that represent diffusing particles such as aSyn monomers. They are geometric objects (spheres and ellipsoids) whose dimensions are in the range of typical intrinsically-disordered proteins.[26] Spherical nanoparticles are constructed by selecting all water beads within a specified radius and tying them together with stiff Hookean springs to create a near-rigid body. Elliptic nanoparticles are created similarly by selecting all water beads within an ellipsoidal volume with given semi-major and semi-minor axes. The simulations place 10, 20, or 30 nanoparticles at the points of a lattice within the solvent region of the box ensuring they do not intersect with the fibril or GFP/linkers if present. The nanoparticles subsequently diffuse freely in the solvent as rigid bodies and are sterically unable to penetrate each other, the GFP tags or the fibril.

### 3) Quantifying the occlusion of the fibril surface by linked GFP tags

The probability that a particle will have its centre of mass in the circular shell from R to R + dR measured from the fibril axis is proportional to the amount of simulation time that it spends in this shell. We have chosen the box size to be sufficiently large that the diffusion of the particles is largely independent. The (unnormalized) histogram of this probability is obtained by summing the number of time-steps in which the particles are in each shell over the simulation time from 2 - 3 10^6^, 3 - 4 10^6^, and 2 - 4 10^6^ time-steps, although we only show a subset of these results for clarity. We discard the first 2 10^6^ steps to allow the system to equilibrate. The histogram is normalised by dividing it by the area of each shell, the number of samples are taken, and the number of particles (see Figure S3 for the limitation due to the simulation box size). It still depends on the length of the fibril and simulation box size. To remove this dependency, we calculate the histogram for the decorated fibril and the corresponding bare fibril (no GFP, no linker). We integrate each histogram over a user-defined region of space around the fibril. The ratio of this integral with the GFP tag to the corresponding integral for the bare fibril defines our measure of the occluding power of the tags. The selection of the precise region of integration is described next.

A dimensionless measure of the fibril surface accessibility is the ratio of the probability of a diffusing particle being within a certain distance of the fibril surface with the GFP tags present to the bare fibril value. This measure depends on the precise region over which the probability is integrated, and requires the lower and upper bounds to be chosen carefully. The lower limit of the integration range is the fibril radius, as the diffusing species are sterically excluded from penetrating the fibril. The upper limit of the integration range is set by the following condition. Far enough from the surface of the fibril, the probability of a particle lying within a given cylindrical shell around the fibril is unaffected by the presence of the GFP tags/linkers. The probability for particles to be in this region should not be included in the occlusion measure as it will overwhelm the signal from the (smaller) region where the GFP tags influence the particle’s motion. The farthest distance at which the GFP tags can sterically interact with the diffusing particles depends on the fibril radius (R_fibril_), linker length (L), and GFP length (4 nm from Yang et al.),[22] R_fibril_ + L + R_GFP_. Thermal fluctuations will reduce this upper limit, so in practise we choose a smaller range by visual inspection of the histograms where the probability distribution is changing most rapidly. We have chosen the upper limit to be 4 nm from the fibril surface. This does not extend to the distance at which the bare fibril probability becomes flat because the signal we are attempting to measure would then be smothered by the probability unaffected by the GFP tag. We have explored the dependence of the occlusion factor when this distance is varied slightly, and our results are not significantly different.

The surface occlusion measure is defined as the ratio of the integrated probability of the nanoparticles’ probability distribution over the predefined range with the GFP tags present to the value for the bare fibril. A value of unity indicates no occlusion, while a value of zero corresponds to the tags completely preventing access to the fibril surface.

## Acknowledgements

The authors express their gratitude to L. Abriata and M. Lopez for critical readings of the manuscript, and S.Thangaraj for the electron microscope image of aSyn fibrils in Figure 1. This study was supported by funding to the Blue Brain Project, a research centre of the École polytechnique fédérale de Lausanne (EPFL), from the Swiss government’s ETH Board of the Swiss Federal Institutes of Technology. The authors gratefully acknowledge computer time provided by the Blue Brain Project and Swiss National Supercomputing Centre. Snapshots and movies of the simulations were produced using the open-source VMD software from the University of Illinois Urbana Champaign (http://www.ks.uiuc.edu/Research/vmd/).[68]

## Author Contributions

**J. C. Shillcock**: Conceptualization, Methodology, Software, Validation, Formal analysis, Writing – original draft, Writing – review and editing, Visualization; **N. Riguet**: Data curation, Writing – review and editing, Visualization; **J. Hastings**: Formal analysis, Data curation, Writing – review and editing, Visualization; **H. A. Lashuel**: Conceptualization, Methodology, Resources, Writing – review and editing, Supervision, Project administration, Funding acquisition

