## Supplemental figures for "Non-monotonic fibril surface occlusion by GFP tags from coarse-grained molecular simulations"

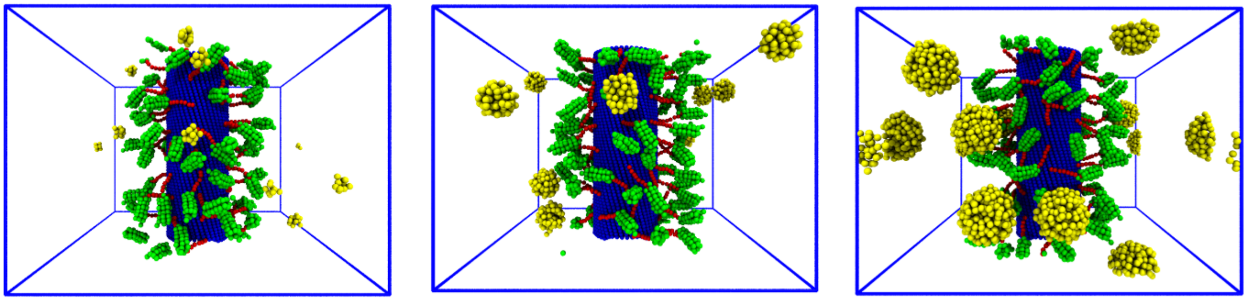

**Figure S1.** Collage of simulation snapshots of 10 spherical particles of radius 1, 2, 3 nm (left to right) around a 10 nm diameter fibril tagged with GFP on 4 nm linkers. The tags are displaced from the fibril's surface and their fluctuations tend to keep the particles farther away from the surface than for the shorter linker. However, when the smallest particles penetrate behind the GFP tags, their escape is hindered by the same steric interactions so that they spend longer at the fibril surface than the larger particles.

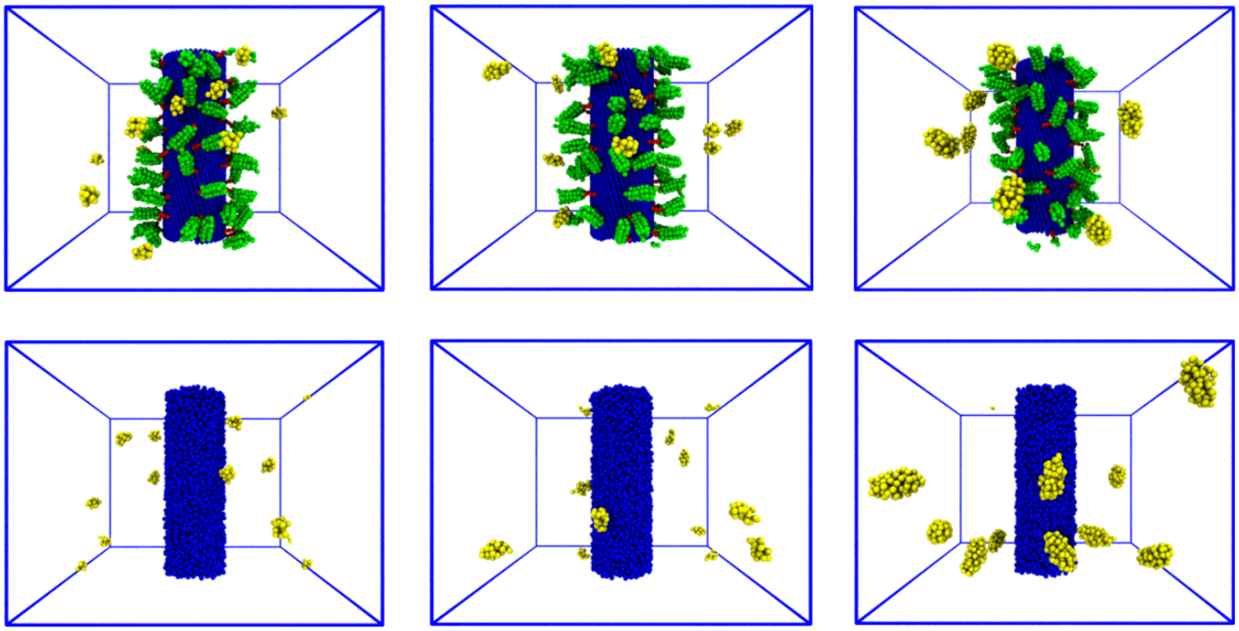

**Figure S2.** Collage of simulation snapshots of 10 ellipsoidal particles of semi-major axes 1.5, 2, 3 nm (left to right) around a 10 nm diameter fibril tagged with GFP on 4 nm linkers (top row) and bare (bottom row). It should be compared to Figure 2 that shows equivalent snapshots for spherical particles. The ellipsoids diffuse around the fibril but are displaced from the fibril's surface by the fluctuations of the tags when present. The actual dimensions of the ellipsoids are: 1.5 x 1 nm, 2 x 1 nm, 3 x 1.5 nm (semi-major x semi-minor axes).

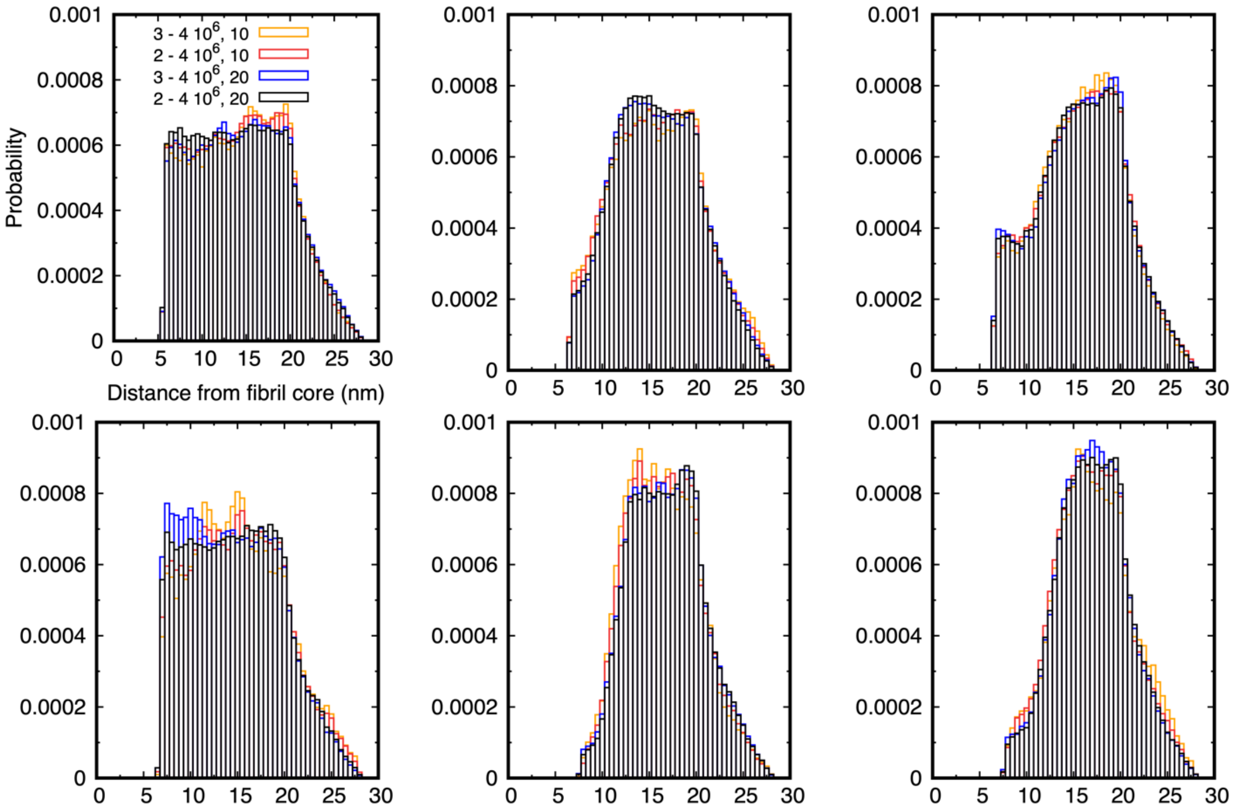


**Figure S3.** (**Above)** Full histograms of the probability for spherical particles (top row 1 nm radius, bottom row 2 nm) to be near the fibril corresponding to those shown in Figure 3. The decrease in the probability beyond 20 nm from the fibril centre occurs because the cylindrical shells protrude beyond the cubic shape of the simulation box. (**Below**) Illustration showing that the radial distribution function of the particles is underestimated when the shell radius exceeds half the width of the simulation box, as the periodic boundary conditions cause the space to wrap around and reduce the number of particles counted in the shell. This does not affect the sampling far from the box boundaries, so the results in Figure 3 are unaffected.

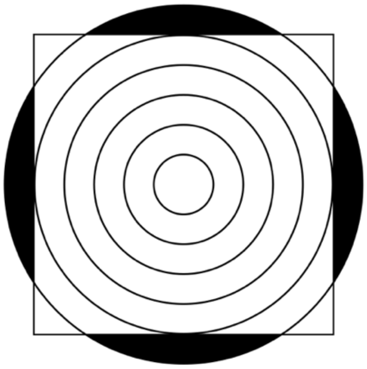

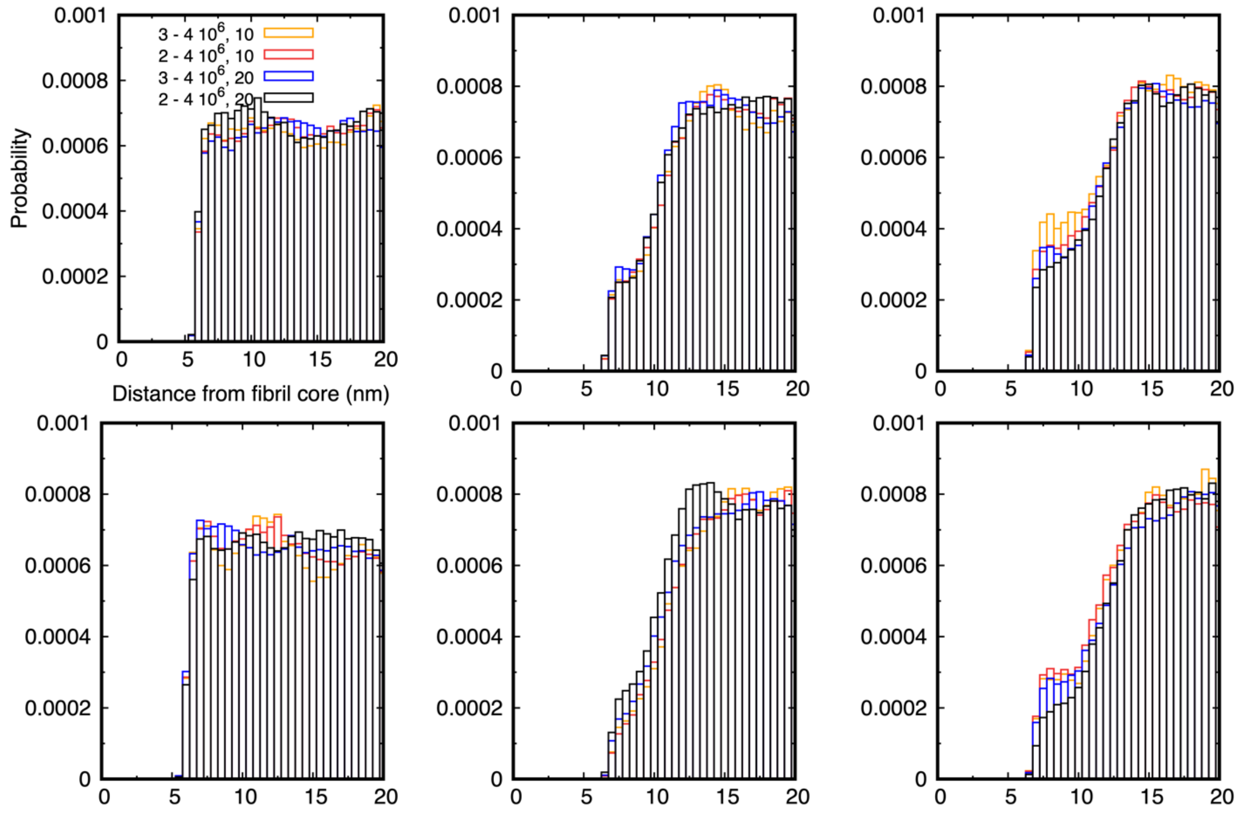


**Figure S4.** Histograms of the probability for 10 and 20 ellipsoidal particles to be distributed around the fibril corresponding to those for spherical particles shown in Figure 3 (top row, 1 nm semi-major axis; bottom row, 2 nm semi-major axis). Left column is for the bare fibril; middle column for GFP tags on 2 nm linkers; and right column for GFP tags on 4 nm linkers. The statistical accuracy of the simulations is indicated by the similarity of the histograms for the different numbers of particles and different sampling periods. In contrast to Figure 3, the reduction in the fibril accessibility on increasing the particle size is less for ellipsoids than spheres and there is a significant enhancement in near-fibril probability for a linker length of 4 nm for both ellipsoid sizes.

Movie SM1 Comparison of the diffusion of 10 spherical nanoparticles with radius 1 nm (left) and 2 nm (right) around a GFP-decorated fibril in which the tags are attached with a 2 nm flexible linker. Note that the solvent particles present in all simulations are invisible for clarity.

Movie SM2 Diffusion of 10 spherical nanoparticles of radius 3 nm around a GFP-decorated fibril with 2 nm linkers showing that the large particles are crowded in the simulation box. The effect is even greater for 20 particles.

Movie SM3 Comparison of 20 spherical nanoparticles of radius 2 nm diffusing around a GFP-decorated fibril with 2 nm linkers (left) and 4 nm linkers (right).

Movie SM4 Comparison of 10 nanoparticles diffusing around a GFP-decorated fibril with 2 nm linkers. The left movie shows spherical particles with a radius of 1 nm while the right movie shows ellipsoidal particles with semi-major axis 1.5 nm and semi-minor axes 1 nm.

Movie SM5 Comparison of 10 nanoparticles diffusing around a GFP-decorated fibril with 2 nm linkers. The left movie shows spherical particles with a radius of 2 nm while the right movie shows ellipsoidal particles with semi-major axis 2 nm and semi-minor axes 1 nm.

Movie SM6 Collage of 10 nanoparticles of radius 1 nm (left), 2 nm (middle), and 4 nm (right) diffusing around an aSyn fibril decorated with GFP on linkers that are 10 nm long. This is more than twice the size of the linkers presented in the paper. The thermal motion of the GFP tags is still able to partially prevent the nanoparticles accessing the fibril surface in spite of the larger free volume behind the GFPs, and the effect increases with increasing nanoparticle size.
